# Metabolite T_1_ relaxation times differ across the adult lifespan

**DOI:** 10.1101/2023.01.06.522927

**Authors:** Saipavitra Murali-Manohar, Aaron T. Gudmundson, Kathleen E. Hupfeld, Helge J. Zöllner, Steve C.N. Hui, Yulu Song, Christopher W. Davies-Jenkins, Tao Gong, Guangbin Wang, Georg Oeltzschner, Richard A.E. Edden

## Abstract

**Purpose:** To investigate the age-dependence of metabolite *T*_1_ relaxation times at 3T in both gray- and white-matter-rich voxels.

**Methods:** This manuscript analyzes publicly available metabolite and metabolite-nulled (single inversion recovery TI = 600 ms) spectra acquired at 3T using PRESS localization. Voxels were placed in posterior cingulate cortex and centrum semiovale in 102 healthy volunteers across 5 decades of life (20s to 60s). All spectra were analyzed in Osprey v2.4.0. To estimate *T*_1_ relaxation times for tNAA_2.0_ and tCr_3.0_, the ratio of modeled metabolite residual amplitudes in the metabolite-nulled spectrum to the full metabolite signal was calculated using the single inversion recovery signal equation. Correlations between *T*_1_ and subject age were evaluated.

**Results:** Spearman correlations revealed that estimated T_1_ relaxation times of tNAA_2.0_ (r_s_ = −0.43; p < 0.001) and tCr_3.0_ (r_s_ = −0.23; p = 0.021) decreased significantly with age in white-matter-rich CSO, and less steeply (and not significantly) for tNAA_2.0_ (r_s_ = −0.15; p = 0.136) and tCr_3.0_ (r_s_ = −0.10; p = 0.319) in gray-matter-rich PCC.

**Conclusion:** The analysis harnessed a large publicly available cross-sectional dataset to test an important hypothesis, that metabolite T_1_ relaxation times change with age. This preliminary study stresses the importance of further work to measure age-normed metabolite T_1_ relaxation times for accurate quantification of metabolite levels in studies of aging.

## Introduction

Understanding the changes in metabolism that occur as the brain ages is an important endeavor in basic and clinical neuroscience. One methodology that can contribute to this effort is proton magnetic resonance spectroscopy ^1^H-MRS), which can measure metabolite levels non-invasively in vivo. MRS-measured metabolite levels have been shown to change in healthy aging^1,2^, and in various diseases^3^. Typically, MRS measurements are acquired with repetition times of 1-5s to improve the SNR efficiency of experiments, re-exciting before the sample is fully relaxed to equilibrium and resulting in signals that are T_1_-weighted (where T_1_ is the longitudinal relaxation time constant). Therefore, in order to infer the concentrations of metabolites from the size of the detected signals, a field-specific T_1_ relaxation correction is required^4–6^.

Current consensus practice in the MRS community^6^ is to perform relaxation correction based upon literature reference values for metabolite relaxation times, because it is not practical to measure metabolite relaxation in each subject. Applying incorrect relaxation correction leads to incorrect quantification of metabolite levels. More crucially, failure to properly account for a range of metabolite relaxation times amongst the study group can lead to incorrect conclusions being drawn. Therefore, in order to perform robust MRS studies of aging, it is important to measure the possible impact of age on relaxation times. Metabolite T_1_S are influenced by local cellular microenvironment, making it plausible that they may change with healthy aging, potentially even representing an interesting biomarker for aging distinct from metabolite levels themselves.

The effect of aging on metabolite T_2_ relaxation times has been previously reported, primarily through measuring metabolite singlets^7–14^. A majority of these studies^8,11–14^ showed a decrease in metabolite T_2_ relaxation times across age; two studies^7,9^ did not find any significant differences between young and aged cohorts; and a third study^10^ showed an increase in metabolite T_2_ with age, using different methodology. T_2_ relaxation times are influenced by changes in the cellular microenvironment and accumulation of iron with age^14,15^.

The impact of age on metabolite T_1_ relaxation times has not been as extensively investigated. Only two studies^7,10^, both performed at 1.5T with small-to-moderate group sizes, have directly compared T_1_ between young and aged groups. The first study^7^ with eight younger subjects (20-30 years) and eight older subjects (60-80 years) found no significant differences in metabolite T_1_ relaxation times. A later study^10^ showed significant decreases in T_1_ relaxation times of the methyl groups of N-acetylaspartate (NAA) and creatine (Cr) with age, including only five older subjects (> 60 years). These differing results have not yet been reconciled, and no attempts have been made to investigate T_1_ across the adult lifespan or at field strengths above 1.5T.

Therefore, this work aims to investigate the age dependence of metabolite T_1_ relaxation times of two major signals in the MR spectrum, namely total N-acetyl aspartate (tNAA_2.0_ at 2.0 ppm) and total creatine (tCr_3.0_ at 3.0 ppm) at 3T. We have previously acquired macromolecular (MM) spectra from posterior cingulate cortex (PCC) and centrum semiovale (CSO) voxels in 102 healthy volunteers across five decades of adult life^16^. These spectra were acquired with pre-inversion at a fixed inversion time (TI) at which metabolite signals are substantially, but not uniformly or perfectly, nulled; the size of residual metabolite signals in these spectra thus directly reports on the rate of T_1_ recovery. Metabolite T_1_ relaxation times were estimated by comparing the modeled residual metabolite amplitudes in these MM spectra to the full metabolite amplitudes in spectra acquired without pre-inversion. Then, the correlation between age and estimated T_1_ relaxation times was evaluated in order to examine the dependence between the two variables.

## Methods

### Study participants

The data analyzed in this study are publicly available on www.nitrc.org and were previously published^16–18^. All acquisitions were performed on a 3T MRI scanner (Ingenia CX, Philips Healthcare, The Netherlands) on 102 healthy volunteers after approval from the local institutional review board (Shandong University School of Medicine). The cohort consisted of approximately equal number of male and female (49 and 53, respectively) participants aged between 20 and 69 years old.

### Data acquisition

Voxels (dimensions: 30 × 26 × 26 mm^3^) were placed in CSO and PCC regions, selected as white-matter (WM)- and gray-matter (GM)-rich brain regions with excellent MRS signal quality. Metabolite-nulled MM spectra were acquired using a single-inversion-recovery PRESS localization sequence^19^ (TR/TE/TI: 2000/30/600 ms) with Philips CHESS^20^ water suppression. Metabolite spectra were also acquired (TR/TE: 2000/30 ms) using PRESS localization and VAPOR^21^ water suppression. Further acquisition details are reported in references^16–18^.

### Data analysis

Data were analyzed using Osprey (v2.4.0)^22^, an open source MRS analysis toolbox, in MATLAB (R2022a). MM spectra were processed as described in Hui et al^16^. In order to address the residual metabolites in the frequency-domain MM spectra, a simulated reduced metabolite basis set including spectra representing total N-acetyl aspartate at 2.0 ppm (tNAA2.0), total creatine at 3.0 ppm (tCr_3.0_), and creatine methylene at 3.9 ppm (CrCH_2_) was used to fit the residual metabolite signals using a linear combination modeling method. A highly flexible spline baseline with 0.1 ppm knot spacing was used to model the MM peaks between 0.5 and 4.2 ppm. In the original manuscript, the fitted metabolite signals were subtracted from the spectra to obtain ‘clean’ MM spectra. In this analysis, the fitted metabolite residual amplitudes themselves (specifically, tNAA_2.0_, tCr_3.0_) are used to measure T_1_. The metabolite spectra (acquired without pre-inversion) were modeled using Osprey’s LCM algorithm^22,23^ and a basis set consisting of 18 metabolites generated by density-matrix simulation in the MRSCloud^24^ toolbox based on FID-A^25^. Metabolites included in the basis set were: ascorbate Asc; aspartate Asp; Cr; negative creatine methylene CrCH2; gammaaminobutyric acid GABA; glycerophosphocholine GPC; glutathione GSH; glutamine Gln; glutamate Glu; myo-inositol ml; lactate Lac; NAA; N-acetylaspartylglutamate NAAG; phosphocholine PCh; phosphocreatine PCr; phosphorylethanolamine PE; scyllo-inositol sl; and taurine Tau. An experimentally acquired MM basis function was included in the basis set in order to model the MM spectrum underlying the metabolite signals, derived from the clean cohort-mean MM spectra^16^. A spline baseline with 0.4 ppm knot spacing was used to account for the background fluctuations in the metabolite spectra. The fitted metabolite amplitudes (tNAA_2.0_, tCr_3.0_) were considered for the T_1_ relaxation time estimation.

### T_1_ estimation

Metabolite residual amplitudes from the MM spectra and metabolite steady state amplitudes from the metabolite spectra were considered for both tNAA_2.0_ and tCr_3.0_.

Theoretically, the signal amplitude in the single inversion recovery experiment is proportional to:

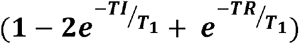

Normalizing this signal to the corresponding steady state signal, the ratio between metabolite signal amplitudes in the two spectra is:

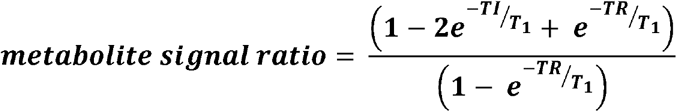

This ratio was calculated between the single-inversion-recovery amplitudes for tNAA_2.0_ and tCr_3.0_ and their steady-state signal amplitudes. With T_1_ being the only unknown, the equation was solved by using nonlinear least squares regression to estimate T_1_ relaxation times of tNAA_2.0_ and tCr_3.0_ in each region. These estimated T_1_ relaxation time values were then plotted against age.

### Statistical Analysis

Statistical analyses were performed using R 4.0.0 (R Core Team, 2021^26^) within RStudio (RStudio Team, 2021^27^). With the exception of T_1_ relaxation time for tCr_3.0_ in PCC, the remaining variables of interest (age, tNAA_2.0_ in PCC and CSO, and tCr_3.0_ in CSO) did not satisfy the normality assumption for Pearson correlations (Shapiro test p < 0.05). Thus, four non-parametric Spearman rank correlations were calculated to examine relationships between estimated T_1_ relaxation times and age for both tNAA_2.0_ and tCr_3.0_ in the CSO and PCC regions separately.

## Results

After visual inspection of the spectra, 4 and 6 datasets from CSO and PCC were excluded respectively, one due to visible ethanol signal in the metabolite spectrum and others due to high lipid contamination. In total 98 and 96 datasets were considered for further analysis. Overall quality of both MM (average NAA linewidth: 6.6 Hz in both CSO and PCC) and metabolite (average NAA linewidth: 6.8 and 7.0 Hz in CSO and PCC respectively) datasets from both CSO and PCC was excellent.

Figure 1 shows an MM spectrum acquired with T_1_ 600 ms and a metabolite spectrum, both from the CSO region of one subject. Negative tNAA_2.0_ and tCr_3.0_ metabolite residuals can be seen in the MM spectrum. The modeled tNAA_2.0_ and tCr_3.0_ components are also shown. The minimal modeling residual shows good fit quality for both MM and metabolite spectra.

**Figure 1:**
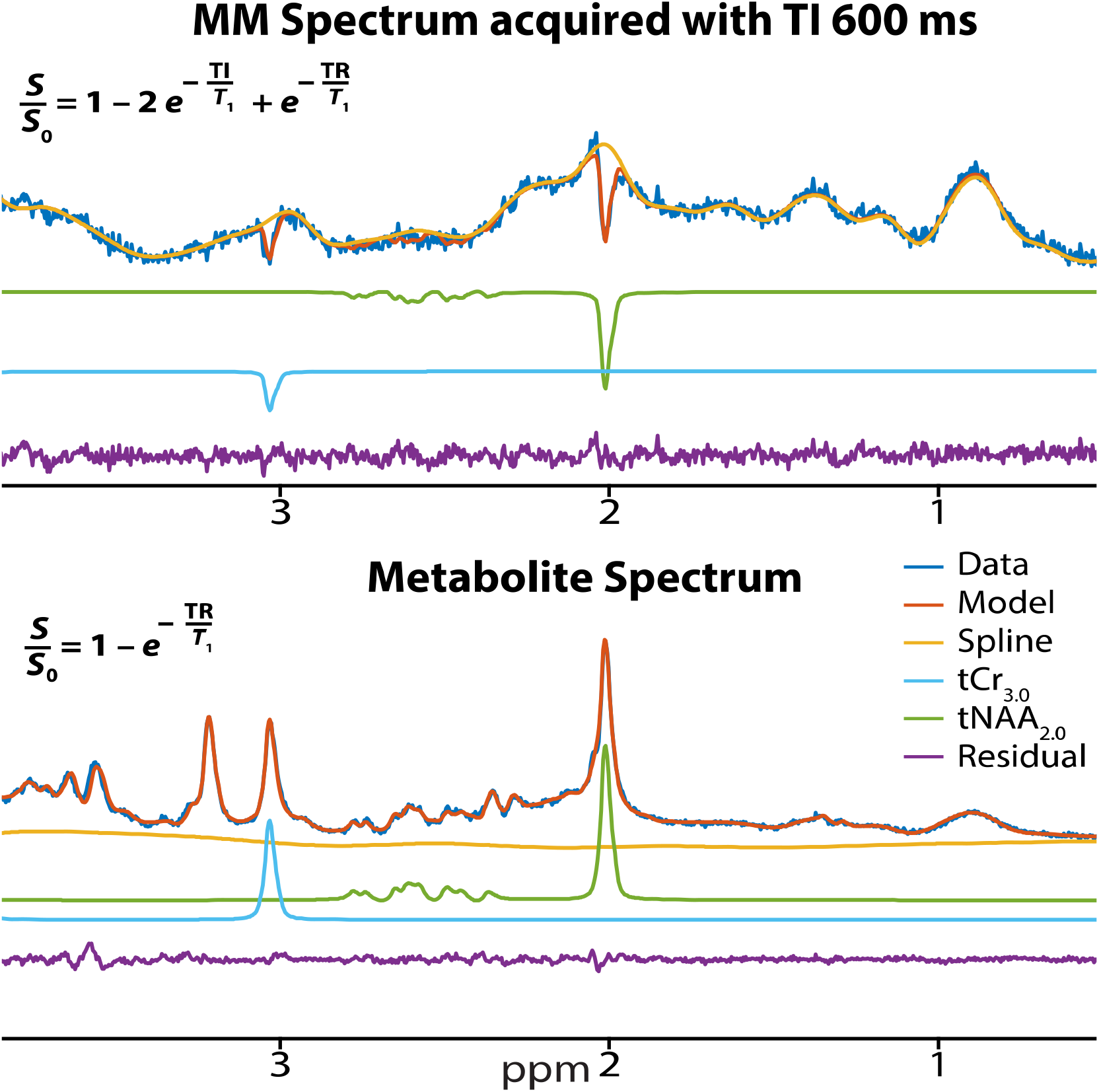
Pre-inverted MM (top) and steady-state metabolite (bottom) spectra and models from a representative subject in CSO. The inlaid equations give the signal amplitudes for the respective spectra.

Plots of estimated metabolite T_1_ relaxation times for tNAA_2.0_ and tCr_3.0_ in PCC and CSO against age are shown in Figure 2. Spearman correlations indicated that older age was significantly correlated with lower T_1_ relaxation times for both tNAA_2.0_ (r_s_ = −0.43; p < 0.001) and tCr_3.0_ (r_s_ = −0.23; p = 0.021) in the WM-rich CSO region. In the GM-rich PCC region, although estimated T_1_ relaxation times were also lower in older age, these relationships did not reach statistical significance for tNAA_2.0_ (r_s_ = −0.15; p = 0.136) or tCr_3.0_ (r_s_ = −0.10; p = 0.319).

**Figure 2:**
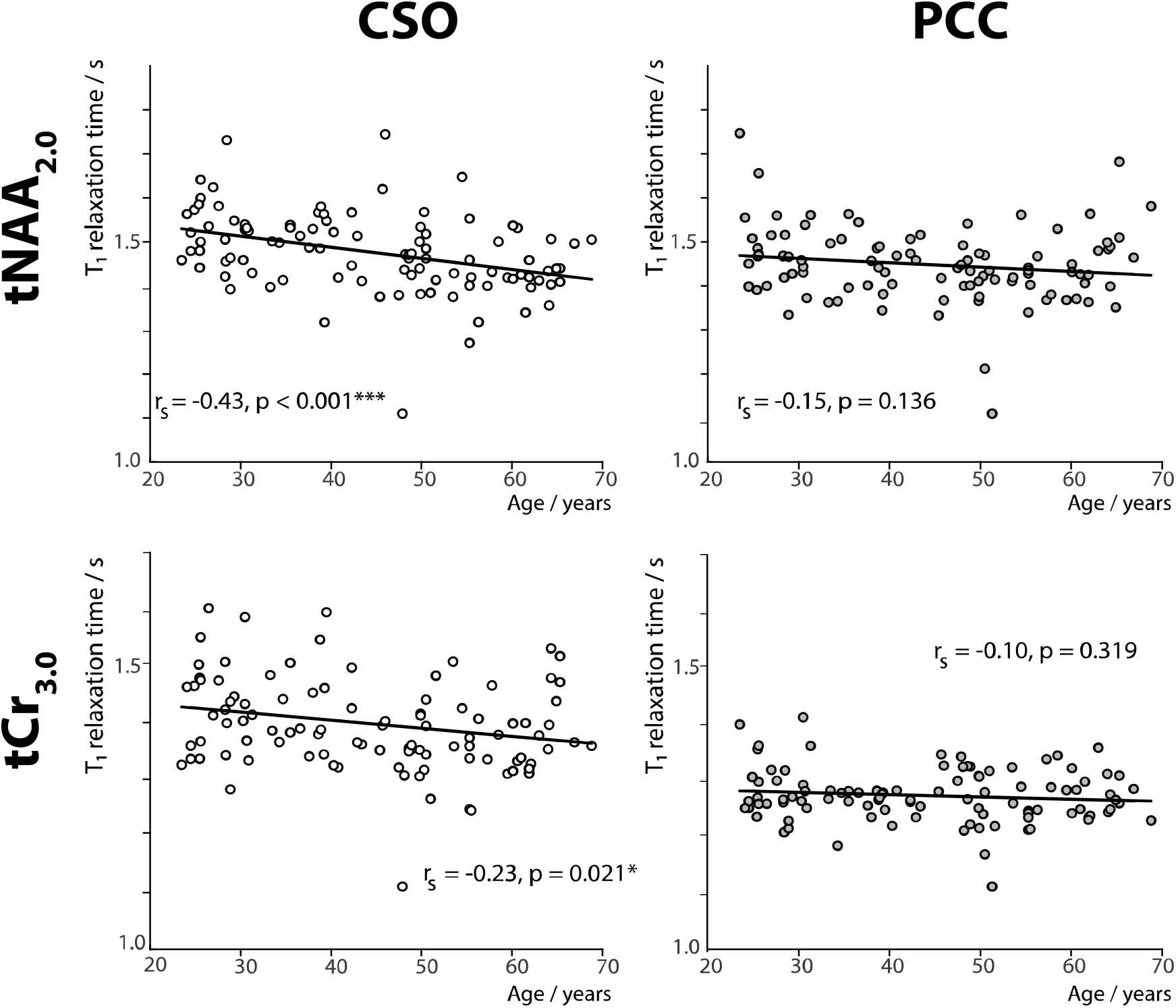
Correlation plots of estimated T_1_ relaxation time for tNAA_2.0_ and tCr_3.0_ against age in CSO and PCC. r_s_ = Spearman correlation coefficient.

## Discussion

We present a novel analysis of a large publicly available cross-sectional dataset (data available on https://www.nitrc.org/projects/mm_mrs/) that was acquired previously^16–18^ from 102 healthy volunteers to characterize the MM and metabolite spectra in aging. Metabolite residual subtraction for MM spectra is implemented in the open-source MRS processing toolbox Osprey according to consensus-recommended^28^ methods. The different T_1_-weighting states of the MM and metabolite spectra can be exploited to estimate metabolite T_1_ relaxation times. Generally, increasing age was associated with decreasing estimated T_1_ relaxation times, but the effect was stronger for tNAA_2.0_ than tCr_3.0_ and stronger in the CSO than the PCC region.

*T_1_* values derived from this two-point measurement should be considered cautiously. The single inversion equation considered for estimating the T_1_ relaxation times of the metabolites assumes perfect inversion. The inversion pulse is an adiabatic pulse applied at 1.9 ppm with a full-width half-maximum bandwidth of 698 Hz, so should perform relatively well throughout the acquired voxel for both NAA_2.0_ and tCr_3.0_ signals. However, the distinction between ‘relatively well’ and perfect inversion may still be meaningful^29^, due to B_1_ inhomogeneity and inherent pulse imperfections. Although the inversion null-point is a TI at which signals are extremely sensitive to T_1_, quantification of the relatively small residual signals (including discrimination by the model between those signals and the MM signals) is challenging. Despite these imperfections, the robust size and age balance of this cohort in comparison to prior work^7,10^ and the lack of prior 3T evidence makes this work a substantial datapoint in the important effort to characterize the relationship between age and longitudinal relaxation times.

Extracting T_1_ relaxation times of tNAA_2.0_ and tCr_3.0_ at age 30 from the trendline in Figure 2 gives values of 1518 ms and 1390 ms respectively in CSO and 1474 ms and 1267 ms respectively in PCC. These values are in the same ballpark as previous literature, although mostly longer than the previous 3T study^30^ (in which T_1_ relaxation times were 1350 and 1240 ms for tNAA_2.0_ and tCr_3.0_ in WM and 1470 and 1460 ms for tNAA_2.0_ and tCr_3.0_ in GM). This prior work measured peak heights manually from the spectra, a less robust approach than LCM, which might have led to errors in the T_1_ relaxation time measurements. Investigating the trend of T_1_ relaxation times of tNAA_2.0_ and tCr_3.0_ across different field strengths, the *T*_1_ values reported in this study are longer than the values reported at l.5T^31^ and mostly shorter than the values reported at 7T^32^. In our results, T_1_ relaxation times were longer in WM than in GM for both signals. This GM-WM T_1_ trend does not agree with the previous work at 3T^30^. However, it matches other prior human studies at different field strengths (1.5T, 7T, and 9.4T) that also show longer metabolite T_1_ values in WM than in GM.

Although not reported here, it is worth noting that the tissue segmentation of both CSO and PCC voxels based upon T_1_-weighted MPRAGE water images does change with age. The GM tissue fraction reduces significantly with age in PCC, and the WM tissue fraction reduces significantly with age in CSO as reported in Hui et al^16^. It is not clear however, whether this represents true tissue volume changes or changes in contrast and segmentation. It is well-known that GM/WM contrast in Tl-weighted images is reduced with aging, making segmentation harder and potentially biasing the voxel fractions away from the major component in each case. Most of the aging literature suggests that loss of WM is the major structural change in healthy aging^34–36^. CSF content in both voxels increased significantly (CSO: R=0.49; PCC: R=0.37) with age^16^. On average, the PCC voxel was 28% GM and 61% WM and the CSO voxel 79% GM and 19 % WM. Our results, i.e. decreasing T_1_ relaxation times with age in the WM-rich CSO and a weaker trend in the GM-rich PCC, may largely be explained by WM changes.

From the slopes and intercepts of the trendlines from Figure 2, the percentage change in the 1 - exp(-TR/T_1_) term for tNAA_2.0_ and tCr_3.0_ in CSO is 8.2% and 5.0%, and in PCC is 3.4% and 1.6% respectively, comparing 20 years of age to 70 years of age. This exponential term is used in correcting the T_1_ weighting when calculating concentration values. Using current analysis methods that assume a single metabolite T_1_ is valid for all ages, this change in T_1_ weighting would be incorrectly interpreted as an increase in metabolite concentration with age and potentially also lead to inconsistent results between aging studies acquired with different parameters.

## Conclusion

In conclusion, this study provides strong evidence of an age-dependence of metabolite T_1_ relaxation times. Estimated T_1_ values decreased with age for tNAA_2.0_ and tCr_3.0_, particularly in the WM-rich CSO. Hence it is suggested that age-specific metabolite T_1_ relaxation times need to be considered in aging studies to measure metabolite concentrations with accuracy. Further focused work is necessary in order to determine accurate T_1_ relaxation times of all observable metabolites across the adult lifespan.

